# Systems modelling of TGF-β/Hippo signalling crosstalk uncovers molecular switches that coordinate YAP transcriptional complexes (submitted to *iScience*)

**DOI:** 10.1101/2022.04.06.487403

**Authors:** Milad Ghomlaghi, Mandy Theocharous, Sung-Young Shin, Eric O’ Neill, Lan K. Nguyen

## Abstract

The Hippo pathway is an evolutionarily conserved signaling network that integrates diverse cues to regulate cell fate and organ homeostasis. The central downstream pathway protein is the transcriptional co-activator Yes-associated protein (YAP). Although capable of inducing gene transcription, YAP cannot bind DNA directly. Instead, it mediates transcriptional activity through interaction with distinct DNA-binding transcriptional factors (TFs), including TEAD, SMAD, and p73, to form active and functionally opposing transcriptional complexes. Growing evidence in mammals demonstrates that YAP has a dual role and can either promote cell proliferation or apoptosis, which underpin its ability to function as both an oncogene or a tumour suppressor depending on the specific context. However, the mechanisms by which YAP coordinates its distinct transcriptional complexes and mediates context-dependent function remain poorly defined. This is in part due to the lack of systems-level studies that can decrypt the complexities of upstream signalling pathways and their crosstalk, which together dictate the transcriptional regulation at the YAP level. Here, we undertake an integrative systems-based approach combining computational network modelling and experimental studies to interrogate the dynamic formation of and transition between the YAP-SMAD and YAP-p73 transcriptional complexes, which control proliferative and apoptotic gene expression, respectively. We developed a new experimentally-validated mathematical model of the TGF-β/Hippo signalling crosstalk and used this model to elucidate dynamic network behaviour. Our integrative studies uncovered previously unknown molecular switches that control the YAP-SMAD/p73 complexes in an on/off, switch-like manner. RASSF1A and ITCH were identified as major regulators of the switches, whereby a graded increase in ITCH expression can trigger YAP to abruptly switch from binding p73 to SMAD, swiftly promoting proliferative gene expression. Further, adjusting the model to reflect cell type-specific protein expression profiles using both in-house and publicly available experimental data enabled us to study the YAP switches under diverse and varied cellular contexts. Overall, our studies provide a new quantitative and systems-level understanding of the dynamic regulation of functionally opposing YAP transcriptional complexes in mammalian cells.

## INTRODUCTION

The Hippo signalling pathway plays critical role in the regulation of tissue homeostasis, organ size, cell proliferation, and apoptosis (1, 4-7). Dysregulation of the pathway in human has been linked to cellular hyperproliferation, invasion and metastasis that are hallmarks of oncogenesis (8). Yes-associated protein (YAP) is a primary transcriptional effector of the Hippo pathway, located downstream of the core pathway kinases MST1/2 and LATS1/2 (9). Although possessing transcriptional activity, YAP cannot bind DNA. Instead, it interacts with a number of DNA-binding transcription factors (TFs), including TEAD, SMAD, RUNX and p73, to form transcriptionally active complexes and to induce target genes’ expression. While it is clear that YAP is an important positive regulator of pro-proliferative and anti-apoptotic genes, growing evidence suggests that in mammals YAP has a dual role, and can either inhibit or promote apoptosis (10). This seems to underpin the ability of YAP to function both as an oncogene or tumour suppressor depending on the biological context (10). However, the molecular mechanisms by which YAP coordinates its context-dependent function remain incompletely understood. Clearly, improved mechanistic understanding of this process may help identify novel strategies targeting Hippo-YAP pathway dysregulation in human cancer.

The dichotomous function of YAP stems in part from its capacity to bind and form complexes with different TFs that have distinct biological functions. For example, members of the TEAD transcriptional factor family are major binding partners of YAP, which upon YAP binding activates target genes involved in cell proliferation (11). YAP also binds SMAD and YAP/SMAD transcriptional complexes to contribute to oncogenic phenotypes (12, 13). On the other hand, YAP can bind to DNA-binding tumor suppressors including RUNXs and p73 to mediate its anti-oncogenic function. RUNX3 interacts directly with TEAD, which markedly reduces TEAD’s DNA-binding ability and disrupts YAP-TEAD mediated oncogenic gene expression in gastric cancer (14). RUNX1 and 3 are also negative regulators of the oncogenic function of YAP in breast cancer (15). To restrict cell growth or respond to stress, the Hippo pathway increases YAP affinity for p73 to promote the transcription of pro-apoptotic genes (16). In this context, we and others have identified the tumour suppressor and scaffold protein RASSF1A as a key positive mediator of YAP-p73 complex formation via signalling through the MST1/2-LATS1/2 kinase cassette (2, 17-19). Despite the increasing evidence supporting that YAP may coordinate its oncogenic or tumor-suppressing function by switching between different binding partners, how this occurs remain poorly defined.

The coordination of distinct YAP-TFs complexes and ensuing cell fates is through signalling pathway crosstalk. We previously showed that crosstalk between the RASSF1A/MST2/LATS and RAF-1/ERK signalling modules orchestrates the decision between cell survival and apoptosis, which is mediated by signalling switches arising from intricate competing protein interactions (1). In addition, the Hippo pathway is engaged in crosstalk with various other pathways, including RAF-1/ERK (1, 20), WNT and PI3K/AKT (21, 22). The TGF-β/SMAD signalling pathway, which plays a critical role in various stages of tumorigenesis, also intimately interplays with Hippo-YAP signalling (3, 23-27). Stimulation by TGF-β triggers dimerisation and activation of the TGF-β Receptors (TGF-βR1/2), which induces phosphorylation of SMAD2/3, complex formation with SMAD4, and translocation to the nucleus to engage DNA-binding partners and transcriptional co-activators for transcription of target genes (28). The nuclear-cytoplasmic shuttling of SMADs is tightly regulated and is mediated through association with YAP (26). In the absence of YAP/ TAZ, SMADs fail to accumulate in the nucleus, and TGF-β mediated transcription is disabled (26). The YAP-SMAD complex also can bind TEAD in the nucleus and promotes TGF-β mediated tumorigenic activity (12, 13). Furthermore, we previously demonstrated that TGF-β/Hippo signalling crosstalk is further mediated by the interaction between TGF-βR1/2 and RASSF1A (3). Upon TGF-β mediated activation of the receptors, RASSF1A is recruited to the plasma membrane (PM) together with the E3 ligase ITCH where it is ubiquitinated and degraded by ITCH. The degradation of RASS1A leads to enhanced formation of the YAP-SMAD complex that promotes a pro-proliferative gene expression program (Fig. 1A). As RASSF1A is a known regulator of YAP-p73 binding, these data suggest that RASSF1A and ITCH may coordinate the assembly of YAP complexes with the different TFs, SMAD and p73. Yet, given the complexity of the TGF-β/Hippo crosstalk network, a systematic understanding of the dynamic regulation of the YAP-SMAD/p73 complexes is lacking.

**Figure 1.**
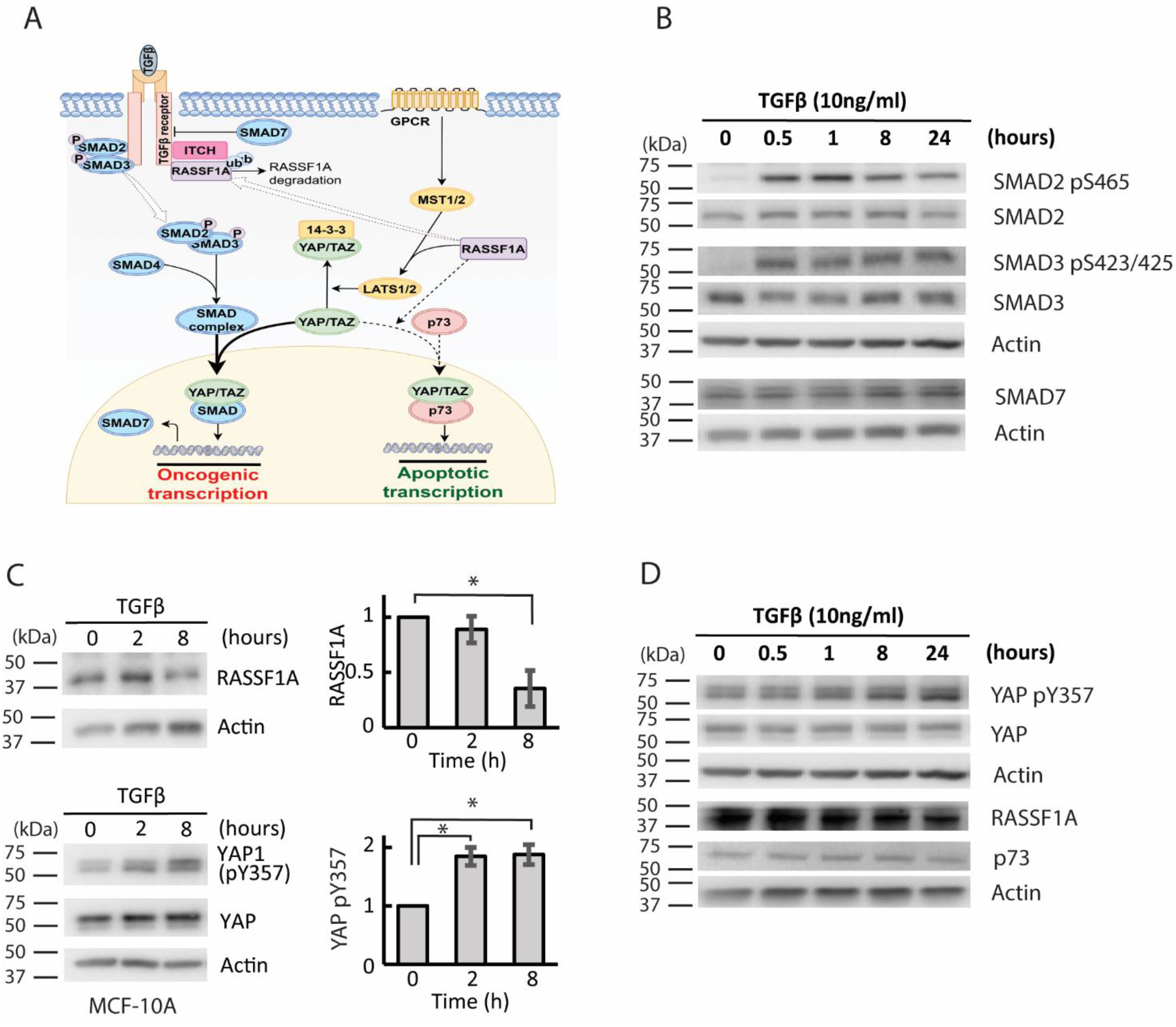
Characterization of the dynamic TGF-β-Hippo pathway signalling crosstalk. **(A)** A schematic diagram displaying the crosstalk between the TGF-β/SMAD and Hippo/RASSF1A/YAP signalling pathways. **(B)** Time course analysis of TGF-β stimulation on the TGF-β/SMAD signalling pathways in the MCF-10A cell line. Expression and activation of signalling proteins in 0 (untreated condition), 0.5, 1, 8 and 24 h post-treatment with 10 ng/ml of TGF-β. Lysates were immunoblotted with antibodies as specified, with actin as a loading control (n=3, p; phosphorylated). **(C)** Time course analysis of TGF-β stimulation on RASSF1A and phosphorylated and total YAP in MCF-10A cells. Expression and activation of signalling proteins in 0 (untreated condition), 0.5, 2, and 8 h post-treatment with 10 ng/ml of TGF-β. Quantification of the western blot data is given on the right panels (* indicates p-value < 0.05, two-tailed student’s t-test, n = 3 biological replicates). **(D)** Time course analysis of TGF- β stimulation (10 ng/ml) on components of the Hippo/RASSF1A signalling axis in MCF-10A cells, as done in (C) but over 24 hours of stimulation.

Here, we undertake an integrative systems-based approach combining computational network modelling and experimental studies to interrogate the dynamic formation of and transition between the YAP transcriptional complexes. We developed a new mechanistic model of the TGF-β/Hippo signalling crosstalk and calibrated this model using time-resolved experimental data. Our integrative studies reveal RASSF1A and ITCH as important regulators of the YAP-p73 and YAP-SMAD complexes. Interestingly, a graded increase in ITCH expression can lead to switch-like response of the YAP-TFs complexes, where YAP abruptly switches from binding p73 to SMAD to promote proliferative gene expression. Model simulations of these switches show the steepness of the switches depends on the protein expression profile within specific cell types. Moreover, increasing ITCH can switch the temporal dynamics of YAP-SMAD complex from a transient to a sustained response following TGF-β stimulation. Using data from DepMap (29), we further found that the essentiality of ITCH in maintaining cell viability significantly correlates with its impact on regulating YAP complexes. These results together with the observed upregulation of ITCH in different tumour types (30, 31) highlights ITCH as a potential therapeutic target for anti-cancer treatment. Overall, our studies provide new insights into the dynamic regulation of functionally opposing transcriptional YAP-TF complexes.

## RESULTS

### Dynamic TGF-β/Hippo signalling pathway crosstalk

Building a dynamic mathematical model of the TGF-β/Hippo signalling crosstalk requires a time-resolved understanding of the interplay between components of the pathways. Thus, to characterize their dynamic crosstalk, we exposed the mammary epithelial MCF-10A cells to TGF-β stimulation and determined the temporal response profiles of the pathways’ components using immunoblot analysis. We observed a rapid and potent increase in the phosphorylation of SMAD2 and 3 following TGF-β treatment as expected, which peaked around 1 hr and gradually reduced toward the latter time points (Fig. 1B, S1A). The transient patterns of pSMAD2/3 are consistent with previous studies and likely due to the known negative feedback mediated by SMAD7 (32, 33). Indeed, we observed a high level of basal SMAD7 already prior to treatment, which increased over time in response to TGF-β and engaged the negative feedback (Fig. 1B). On the other hand, TGF-β treatment led to a significant temporal reduction of RASSF1A expression as found previously (Fig. 1C and S1B), and confirms the inhibitory effect of TGF- β stimulation on RASSF1A (3). Interestingly, we found that TGF-β treatment induced a significant increase in the levels of phosphorylated YAP at tyrosine 357 (pY357), which was double the pre-stimulated level at 2 hr and became sustained at 8 hr post-stimulation (Fig. 1C and S1C). This trend was further corroborated in a longer time-course over 24 hrs of TGF-β treatment (Fig. 1D). YAP phosphorylation at Y357 has been shown to stabilize YAP and induce a stronger affinity for p73 (34). YAP pY357 is mediated by the non-receptor tyrosine kinase c-Abl (34-36), known to be regulated by TGF-β signalling (37). In contrast to YAP pY357, we did not observe significant changes in total YAP or phosphorylated YAP at serine 127 (pS127) after TGF-β treatment (Fig. S1D). Moreover, TGF-β also did not affect the expression of p73 (Fig. 1D, S1E). Together, these results elucidate the dynamics of TGF-β/Hipo-YAP signalling crosstalk downstream of TGF-β stimulation and provide useful time-resolved data for model building. Importantly, our data indicate TGF-β not only triggers SMAD signaling but also potentiates YAP-p73 association through enhanced YAP pY357 (Fig. 1C, S1C), the latter constitutes a novel crosstalk point between the TGF-β and Hippo-YAP pathways.

### A mechanistic model of TGF-β/Hippo-YAP signalling crosstalk

While the evidence for TGF-β/Hippo-YAP crosstalk is clear, how YAP integrates the upstream signals and complex regulatory mechanisms to induce robust transcriptional decisions remains poorly defined. As a basis toward addressing this, we set out to construct a new mathematical model of the signalling network that is aimed to encapsulate the complex crosstalk and regulatory mechanisms under a unified quantitative framework.

#### Model inputs, outputs and reactions

A reaction schematic of the new model is given in Figure 2A, which displays and enumerates the reactions included in the model. The model reactions are described in detail in Table S1. The model’s main input is TGF-β ligand. Briefly, upon TGF-β stimulation, the ligand binds to and activates the TGF-β receptor (reaction R2, Fig. 2A and Table S1), which is subject to protein turnover via synthesis and degradation (R1). Active TGF-β receptor phosphorylates the transcriptional factors SMAD2/3, inducing their complex formation. Since our data show SMAD2 and 3 exhibit similar response dynamics following TGF-β treatment (Fig. 1B and S2B), for simplicity we consider SMAD2 as the representative protein of the SMAD complex (R3) (38, 39). Phosphorylated SMAD2 then forms complexes with YAP and translocates to the nucleus to initiate YAP-SMAD’s target gene transcription (R10) (26). In the nucleus, the YAP-SMAD complex can also associate with TEAD to form ternary complexes. However, in response to stimulation by TGF-β it is likely that SMAD, rather than TEAD, plays a dominant role in driving YAP-mediated proliferative gene expression, allowing us to exclude TEAD from our model for simplicity. In addition, while TGF-β-mediated transcription is highly dependent on YAP (40), it is likely that phosphorylated SMAD2 on its own is also transcriptionally active and thus we assume unbound pSMAD2 can also induce transcription, with a relatively slower rate compared to the YAP-SMAD complex (40). SMAD7 is a target gene of SMAD2/3 (R14), which in turn negatively inhibits TGF-β receptor through receptor ubiquitination and degradation, generating negative feedback (R4) (41).

**Figure 2.**
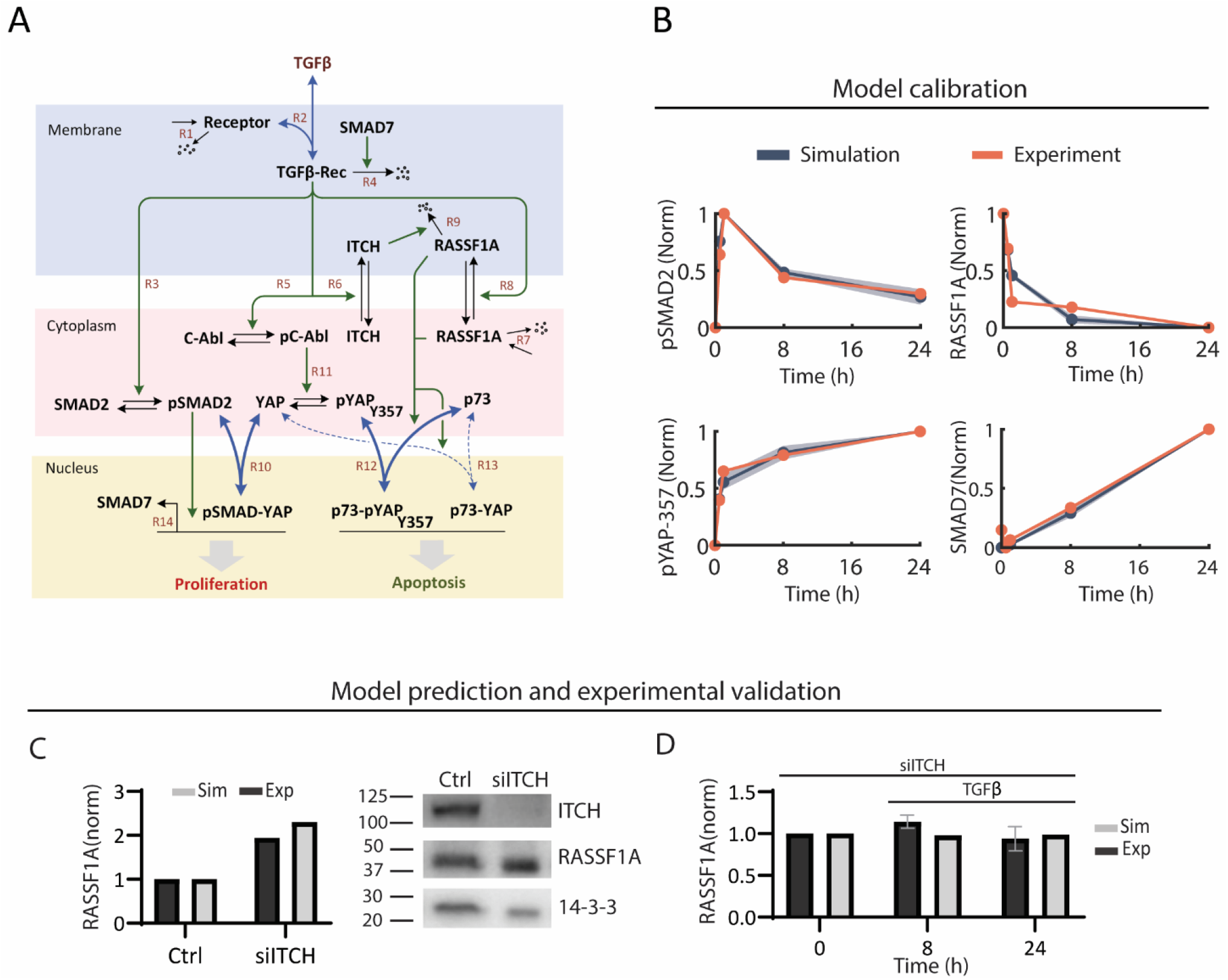
Construction, calibration and validation of a new mathematical model of the TGF-β-Hippo signalling crosstalk network. **(A)** A reaction schematic diagram displaying the biochemical reactions included in the mathematical model of the pathway crosstalk. Detailed description of the model is given in the mai text. **(B)** Time-course model simulations using the best-fitted parameter sets (blue curves) in comparison with the quantified experimental data (brown curves) from Figure 1, showing strong concordance between simulation and data. A total of 11 best-fitted parameter sets obtained from model calibration were used for the simulations (solid blue line demonstrates the mean simulated curve, and shaded area shows standard deviation). **(C, D)** Independent validation of the model. **(C)** Model simulation results predicting upregulation of RASSF1A expression after siRNA-mediated knockdown of ITCH (left), as compared to experimental data determined using western blot (right). Sim = simulation, Exp = experiment. **(D)** Effect of TGF-β stimulation on RASSF1A level under ITCH knockdown condition. Simulations predict ITCH knockdown prevents RASSF1A degradation after TGF-β stimulation, which is confirmed by experimental data. Here, MCF-10A cells where ITCH has been knocked down were stimulated with TGF-β (10ng/ml) and RASSF1A protein expression was measured by western blot.

On the Hippo signalling side, RASSF1A promotes YAP-p73 binding through activation of the kinases MST2/LATS1 (R13 and R14) (1, 2). The potent TGFβ-induced YAP phosphorylation at pY357 (Fig. 1C) suggests this phosphorylation is highly dynamic and tightly regulated, prompting the need to explicitly include YAP-pY357 in the model (R12). YAP phosphorylation at pY357 is catalysed by the kinase c-Abl and can be dephosphorylated by implicit phosphatases to revert to the unphosphorylated form (R11). Since YAP-pY357 binds more favorably to p73 than unphosphorylated YAP (35), we assumed a relatively smaller dissociation constant for reaction R12 compared to R13, and both are stimulated by RASSF1A. We previously demonstrated that following TGF-β stimulation, the TGF-β receptor recruits both RASSF1A and the E3 ligase ITCH to the PM where ITCH degrades RASSF1A (Fig. 1B) (3). Accordingly, the model incorporates this crosstalk mechanism by including the translocation of RASSF1A and ITCH from the cytoplasm to the PM (R8 and R6, Fig. 2A) and ITCH-induced RASSF1A degradation at the PM (R9).

In addition to SMAD and TEAD, YAP can also bind the transcriptional factor RUNX. In normal conditions, RUNX can form complexes with YAP to promote ITCH’s transcriptional expression and p73 degradation in several cell lines (42, 43). However, our experimental data in MCF-10A indicates the levels of both ITCH and p73 were not affected over time by TGF-β treatment (Fig. S2A and S1E), suggesting the YAP-RUNX complex is not regulating ITCH and p73 in this setting. This allowed us to neglect YAP-RUNX binding from our model. Similarly, while LATS1/2 phosphorylate YAP at S127 and retain it in the cytoplasm, our data shows pYAP-S127 levels remained constant (Fig. S1D) after TGF-β stimulation and thus the phosphorylation of YAP by LATS1/2 can be excluded from our model. Together, these experimentally driven assumptions enabled us to simplify the model while keeping the most essential interactions that are critical for governing the network dynamics.

#### Model implementation, calibration and validation

The new model was implemented using ordinary differential equations (ODEs), where the rate of change in the levels of the model species are formulated from the relevant reaction rates that in turn are described using appropriate kinetic laws. The complete list of model ODEs and reaction rates are given in Supplementary Tables S1-3.

To provide the model with context specificity and predictive power, we performed model calibration using the experimental time-course data from MCF-10A cells (Fig. 1B and D). This essentially involves the estimation of unknown model parameters such that model simulations recapitulate the data. Parameter estimation was undertaken in Matlab using a genetic algorithm-based optimization procedure, which was repeated 500 times to identify as many best-fitted parameter sets as possible (see Material and Methods and SI Text for details). This is because performing model simulations using multiple best-fitted sets prevents possible biases associated with any single best-fitted set, thus boosting model prediction confidence. Fig. 2B shows that model simulations using the obtained best-fitted parameter sets reproduce the experimental data well, evidenced by the strong concordance between the simulated and data curves (Fig. 2B).

### Model simulation correctly predicts signalling effect of ITCH knockdown

To further establish the validity of our model, we performed additional model validation using independent experimental data. First, the model simulation predicts that *in silico* knockdown of ITCH upregulates RASSF1A level in the absence of TGF-β stimulation (Fig. 2C). To validate this we knocked down (KD) ITCH using siRNA (Material and Methods) and measured RASSF1A expression in MCF-10A cells using western blot. Consistent with model prediction, ITCH silencing increases RASSF1A concentration by ∼2-folds (Fig. 2C).

Next, we simulated RASSF1A dynamics in response to TGF-β stimulation under ITCH KD condition, which shows no significant change in RASSF1A concentration at 8 and 24 hours after TGF-β stimulation. We tested this experimentally by knocking down ITCH and stimulating MCF-10A cells with TGF-β (Fig. 2D). Consistent with the *in silico* prediction, there were no significant changes in the level of RASSF1A under ITCH KD condition. Together, these independent model validation supports the validity and predictive power of the model.

### RASSF1A and ITCH coordinate switch-like transition between YAP transcriptional complexes

RASSF1A is known to enhance the affinity of YAP for p73 (44). However, how the complex formation of YAP with distinct transcriptional factors, such as p73 and SMAD, is coordinated remains unclear. To address this, we investigate *in silico* the temporal and steady-state responses of YAP-p73 and YAP-SMAD complexes under various network perturbations. First, we performed model simulations where we increasingly varied the synthesis rate of RASSF1A over a wide range and simulated the effect of such perturbation on the levels of YAP-p73 and YAP-SMAD complexes at different time points over 48 hrs of TGF-β stimulation (Fig. 3A, B).

**Figure 3.**
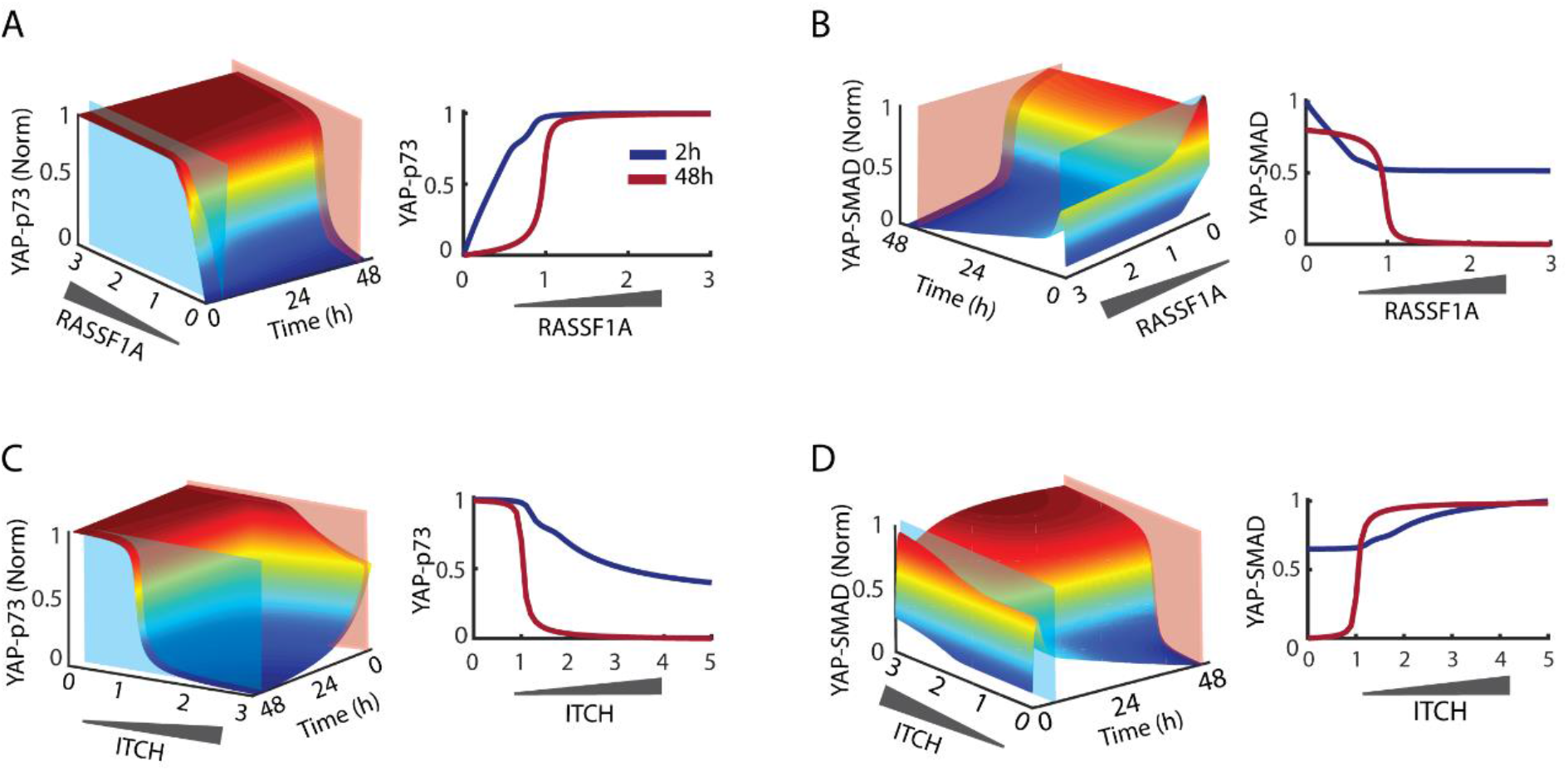
Molecular switches coordinate YAP-SMAD and YAP-p73 transcriptional complexes. **(A-D)** 3-Dimendional (3-D) time-depedent model simulations showing the changes in temporal profiles of the YAP-SMAD and YAP-p73 complexes in response to TGF- β stimulation, for increasing levels of RASSF1A (A-B) or ITCH (C-D). To perturb RASSF1A and ITCH levels *in silico*, the synthesis rate of RASSF1A or the total concentration of ITCH in the model was increased from null to three folds higher than the nominal values, in order to generate a wide range of varying concentrations for each protein. Model simulations predict the presence of abrupt switch-like response of the YAP-TF complexes. For each panel, a 2D simulation snapshot of the RASSF1A-(A,B) or ITCH-(C,D) mediated dose-response curves is presented on the right, in response to early (2h, blue) or late (48h, red) TGF-β stimulation. Simulations show the transition between the YAP-TF complexes are more linear at 2 hours after TGFβ stimulation but become sigmoidal (switch-like) at 48 hours following stimulation.

Model simulation and analysis revealed non-intuitive dynamic properties of the TGF-β/Hippo interaction. At all the simulated time points, increasing RASSF1A induces enhanced YAP-p73 complex formation but inhibits the formation of the YAP-SMAD complex, consistent with the idea that SMAD and p73 compete for YAP binding. Interestingly, while the graded RASSF1A overexpression led to a linear-saturating increase of the YAP-p73 complex at 2 hrs, the response becomes abrupt and displays a switch-like pattern at the late time points, e.g. 48 hrs post-TGF-β stimulation (Fig. 3A). Similar time-dependent input-output effects were observed for the YAP-SMAD complex, but in the opposite direction (Fig. 3A). The identified switches mediate a threshold-gated control governed by RASSF1A, where increasing RASSF1A at a low level has minor effects on YAP-p73/SMAD binding, however, further increase of RASSF1A beyond a certain threshold triggers a dramatic increase in YAP-p73 accompanied by a significant reduction of YAP-SMAD complex.

Pefani *et al*. conducted an experiment following a similar setup using U2OS cells that express RASSF1A under a doxycycline-inducible promoter (3). Using this system RASSF1A expression was gradually increased in the cells and the levels of YAP-SMAD and YAP-p73 binding were measured 2 hrs after TGFβ stimulation (Fig. S3A). The data show that increasing RASSF1A expression suppressed YAP-SMAD binding and promoted YAP-p73 formation. This data validates our model prediction of RASSF1A-mediated transition between SMAD-YAP and YAP-p73 complexes 2 hrs after TGFβ stimulation (Fig. 3A, B).

Upon TGF-β stimulation, both ITCH and RASSF1A are recruited to the PM by TGF-βR where ITCH degrades RASSF1A (3). This led us to hypothesize that ITCH may also regulate the formation and dynamics of the YAP-TFs complexes. To examine this, we simulated the temporal response of YAP complexes over 48 hrs following TGFβ treatment at different expression levels of ITCH, ranging from low to high. Strikingly, increasing ITCH results in a strong switch-on in the formation of the YAP-SMAD complex, accompanied by an abrupt switch-off in YAP-p73 association at the late time points (e.g. 48 hrs). As seen for RASSF1A, these ITCH-mediated switches are not observed at the early TGF-β treatment.

In addition, simulations show that increasing ITCH level switches YAP-SMAD response dynamics from a transient to a sustained pattern. When the level of ITCH is low, TGF-β stimulation leads to an overshoot in YAP-SMAD dynamics that was subsequently attenuated to approximately basal level (Fig. S3B). In contrast, when ITCH level is high TGF-β stimulation results in an increase in YAP-SMAD concentration that remains sustained over time (Fig. S3B). Taken together, these computational analyses suggest ITCH as a critical mediator of the levels and dynamic profiles of the YAP transcriptional complexes.

To understand the network features that underpin the switch-like behaviours, we interrogated the underlying network structure. Switch-like transitions can occur under conditions where post-translational modifications alter protein affinities for binding partners (1). Here, phosphorylated YAP binds p73 and RASSF1A regulates the binding affinity between phosphorylated YAP and p73. Considering that ITCH modulates RASSF1A concentration, at the low level of ITCH there is a high RASSF1A concertation which increases pYAP-p73 binding affinity. However, overexpression of ITCH diminishes RASSF1A and reduces pYAP-p73 complex formation.

### Governing factors of the RASSF1A/ITCH-mediated YAP switches

Here, we aim to determine the governing factors that control the behaviour of the identified RASSF1A/ITCH-mediated YAP switches. While a systematic analysis of this type is challenging experimentally, it is computationally feasible using our model. To this end, we performed model-based sensitivity analysis where we systematically perturbed each model kinetic parameter (10-fold up or down relative to the nominal values) and assessed the impact on the switches’ behaviour. The kinetic parameters generally represent the strength of the network interactions they describe. To assess the change in the YAP switches quantitatively, we characterized a number of general metrics of a switch-like response by fitting the simulated curve to a custom function defined in Figure 4A. The parameter *Km* indicates the threshold (of the input, i.e. ITCH/RASS1A) at which the curve switches on (or off), while *H* represents the steepness of the switch. The higher value of *H* and *K*_*m*_ indicate the switch is steeper and has a higher switching threshold, respectively (Fig. 4B).

**Figure 4.**
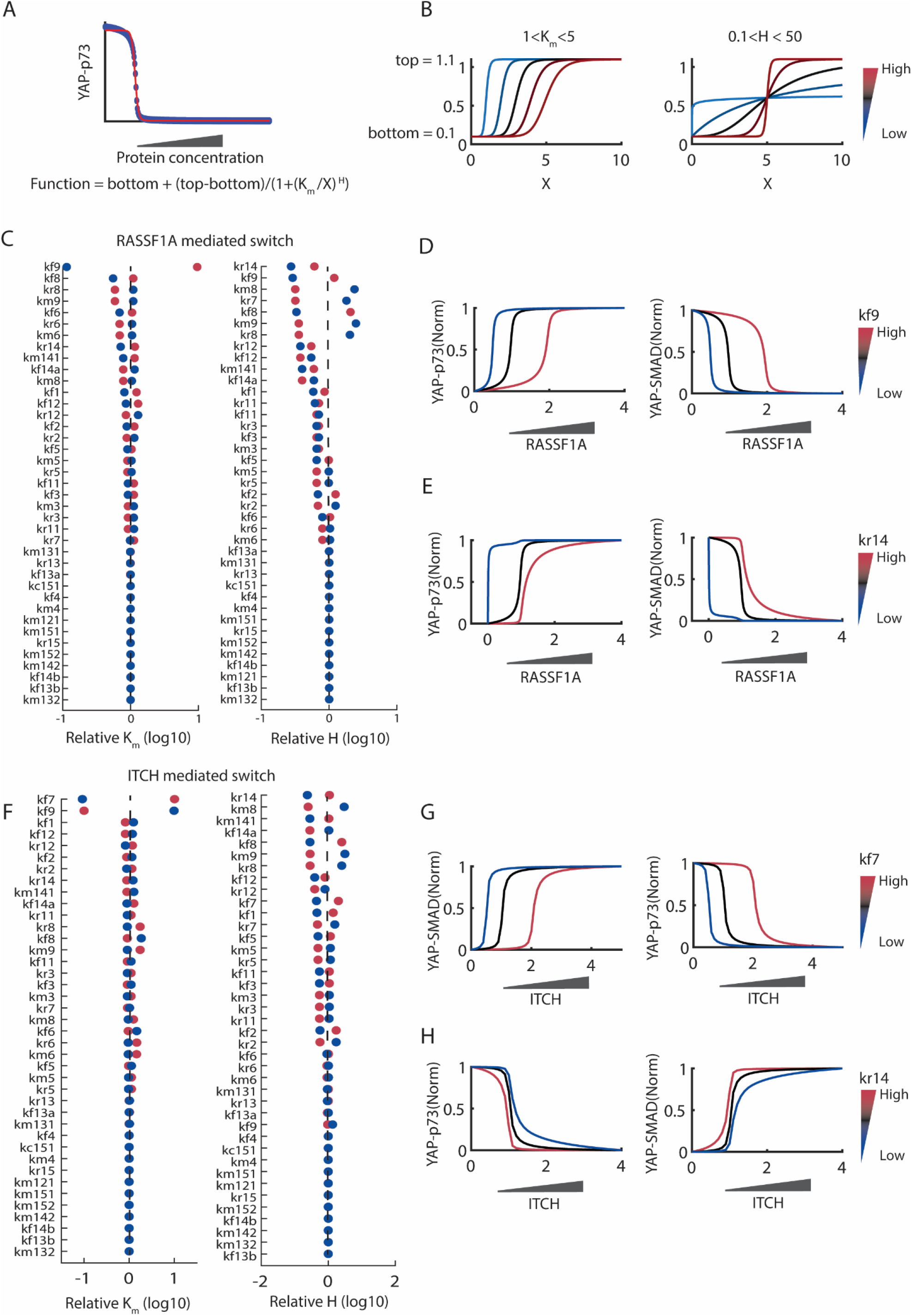
Identification of network factors governing the YAP switches. **(A)** A schematic diagram depicting how the simulated YAP-TF response curves were fitted with a Hill function, in order to quantify the switches’ salient properties, i.e. the switch steepness (parameter *H*) and threshold (parameter *Km*). **(B)** Simulations that illustrate indeed: increasing *K*_*m*_ results in a higher switching threshold, evidenced by the curve shifting to the right; while increasing *H* results in a steeper, more abrupt switch. **(C)** Sensitivity analysis results for the RASSF1A-mediated switch. Each displayed kinetic parameter was systematically perturbed (i.e. 10-fold up or down from the nominal values), and the effects on the switch steepness and threshold were computed. Red dots indicate the effects resulting from increasing the parameters, whereas blue dots indicate the effects from decreasing the parameters. **(D)** Simulation of the YAP-TF complexes in response to a gradual increase in RASSF1A level, at increasing values of the parameter *k*_*f9*_. **(E)** Simulation of the YAP-TF complexes in response to a gradual increase in RASSF1A level, at increasing values of the parameter *k*_*r14*_. **(F)** Sensitivity analysis results for the ITCH-mediated switch. Notations are the same as in (C). **(G)** Simulation of the YAP-TF complexes in response to a gradual increase in ITCH level, at increasing values of the parameter *k*_*f7*_. **(H)** Simulation of the YAP-TF complexes in response to a gradual increase in ITCH level, at increasing values of the parameter *k*_*r14*_.

Figure 4C displays the sensitivity analysis results where the kinetic parameters are ranked according to their impact on the RASSF1A-induced switches. About one-third of the considered parameters had no effects on both the switches’ threshold or steepness (Fig. 4C). The degradation rate of RASSF1A (*k*_*f9*_) exerts the strongest influence on the switching threshold, where increasing *k*_*f9*_ markedly raises the level of RASSF1A required to switch YAP-p73 on and YAP-SMAD off (Fig. 4D). Following *k*_*f9*_, the rates governing RASSF1A translocation to and from the PM (*k*_*f8*_/*k*_*r8*_) are the next most influential parameters in controlling the threshold (Fig. 4C). Interestingly, we found that the dissociation rate of YAP-p73 (*k*_*r14*_) exerts the strongest impact on the steepness of both RASSF1A-and ITCH-mediated YAP switches (Fig. 4C, F). Blocking this rate converts the YAP-p73/SMAD responses from being switch-like to a more smooth pattern (Fig. 4E, H).

In addition, the results show that the synthesis rate of RASSF1A (*k*_*f7*_) has the strongest effect on the threshold of the ITCH-mediated YAP switches. Raising RASSF1A’s synthesis rate potently shifts the YAP-SMAD on-switch and YAP-p73 off-switch to the right (Fig. 4G), meaning higher ITCH levels are required to trigger the switching. This is because as *k*_*f7*_ increases, higher levels of ITCH are needed to degrade the more abundant RASSF1A to be under a critical threshold necessary for the switch between YAP-SMAD and YAP-p73 to occur (Fig. S4A, dashed line). Consistent with this data, increasing RASSF1A’s degradation (rate *k*_*f9*_) has the opposite effect (Fig. 4F). These results together suggest that when RASSF1A is overexpressed, cells require a higher abundance of ITCH to switch on YAP-SMAD and ensuing proliferative transcription program. On the other hand, in tumour where RASSF1A is downregulated we would expect cells do not need to express high levels of ITCH compared to those where RASSF1A is not downregulated. Indeed, analyzing the co-expression profiles of RASSF1 and ITCH across large cohorts of breast and pan-cancer patients shows that patients with down-regulated RASSF1 rarely exhibit overexpression of ITCH compared to those with normal or upregulated RASSF1 (Fig. S4B), consistent with our prediction.

### Interrogation of the YAP switches in different cell types

Our integrative studies have revealed the switch-like behaviour of the YAP-p73/SMAD complexes using the MCF-10A cell line so far. Here, we ask whether these switches may be a common feature in other cellular contexts by examining other cell types. Previously, we determined that cell-to-cell variation in protein expression levels is a major source of signalling response heterogeneity (45). This led us to hypothesize that the YAP switches’ behaviour may be dictated by the protein expression profile across specific cancer cell types. To test this, we first determined the basal protein expression of the network nodes in three additional breast cancer cell lines Hela, MDA-MB-231 and MDA-MB-468 using antibodies-based immunoblotting (Fig. 5A). Data quantification and normalization relative to MCF-10A cells demonstrate marked cell-to-cell protein expression variability (Fig. 5B). Notably, MDA-MB-468 expresses highly abundant p73, the level of which is ∼15, 5 and 2 folds higher than in MCF-10A, Hela and MDA-MB-231, respectively. In addition, MDA-MB-231 and MDA-MB-468 express comparable levels of YAP, which are ∼2-3 folds higher than in MCF-10A and Hela cells. On the other hand, Hela and MDA-MB-231 display the highest basal levels of SMAD2, while all four cell lines exhibit similar expression of ITCH (Fig. 5B). These data together highlight marked variation in the abundance of the network components, which may result in differential network dynamic behaviours between the cell types.

**Figure 5.**
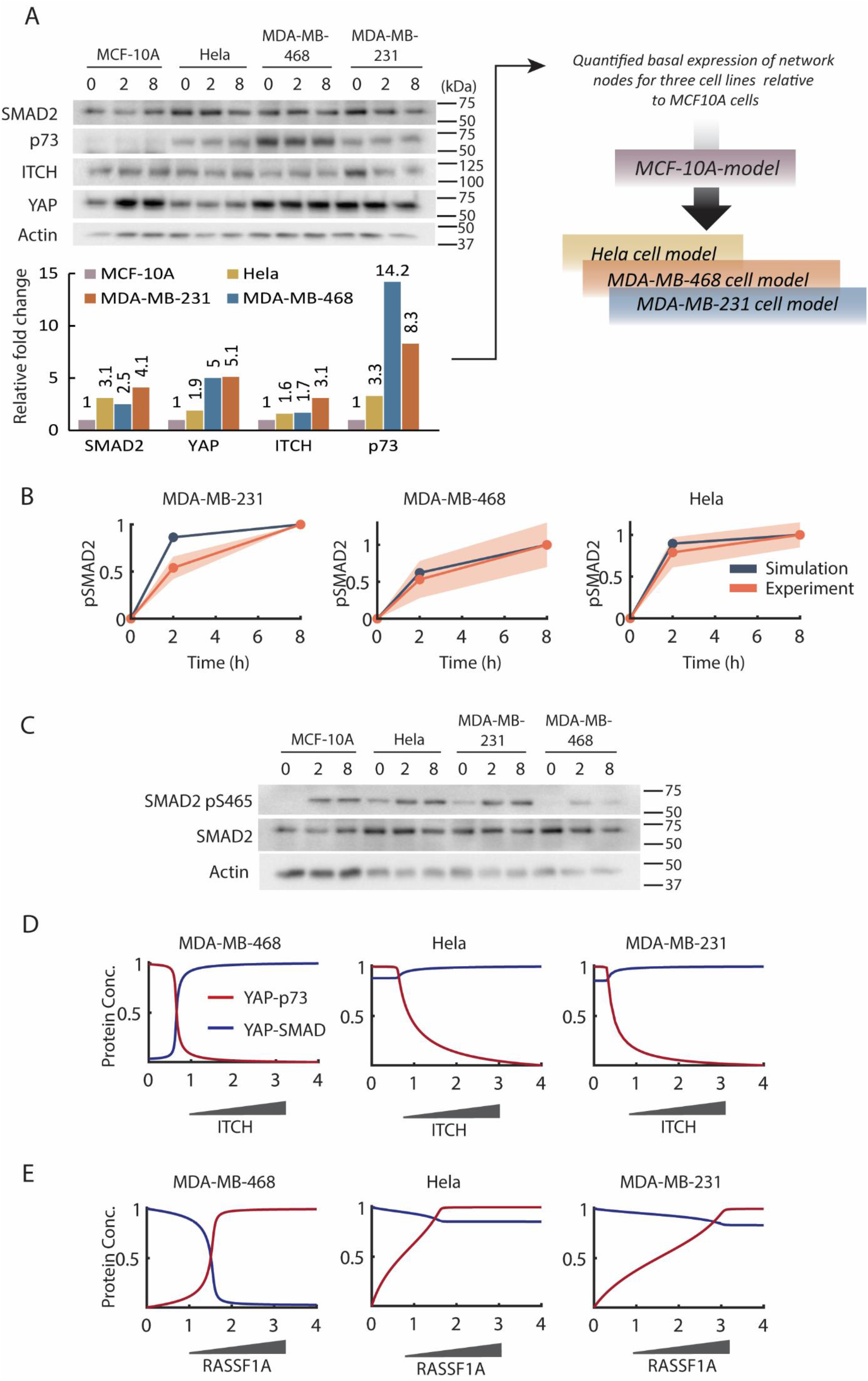
Cell type-specific models predict context dependent signalling activation and switch behaviours. **(A)** Comparative analysis of the basal expression of various network components and their reponse to 8 hours of TGF-β (10ng/ml) across MCF-10A and different breast cancer cell types (Hela, MDA-MB-468 and MDA-MB-231) using western blot (upper panel). Quantification of the data (above) showing the total basal expression of the network components in MDA-MB-468, MDA-MB-231 and Hela cells relative to that in MCF-10A cells (lower pannel). A schematic showing how the relative protein expression data were used to customize the MCF-10A model to generate models for the MDA-MB-468, MDA-MB-231 and Hela cell lines (right pannel). **(B)** Time-course model prediction of phosphorylated SMAD2 in response to TGF-β stimulation in specific cell types using the models constructed in (A), and validates against corresponding experimental data quantified from (C). The orrange line indicates mean and the shaded area indicates standard deviation of three experimental replicates (see also Fig. S5). **(C)** Comparative time-course analysis of TGF-β stimulation (10 ng/ml) on phosphorylated and total levels of SMAD2 in four different cell lines. **(D-E)** Model simulations of the formation and transition between the YAP-TF complexes using the cell type-specific models, which demonstrate highly context-dependent switching behaviour. To perturb ITCH and RASSF1A levels *in silico*, the total concentration of ITCH and synthesis rate of RASSF1A were perturbed from zero to four fold higher than their nominal values.

To investigate this, we utilized the relative expression fold-changes obtained above as inputs to tailor the MCF-10A model for each of the three additional cell lines by adjusting the concentration of the relevant model species accordingly (Fig. 5C). For example, since YAP is 5-fold more abundant in MDA-MB-231 compared to MCF-10A, the level of YAP in the model describing MDA-MB-231 was increased by 5 folds. We have successfully employed a similar model customization strategy in previous studies (1, 46). As a result, we obtained three new models that are specific to Hela, MDA-MB-231 and MDA-MB-468. To test the validity of these models using independent data not used in the model customization process, we simulated the dynamic response of phosphorylated SMAD2 to TGF-β stimulation using each model, and undertook corresponding time-course experiments to confirm model predictions (Fig. 5D,E). Overall, the simulations are consistent with the qualitative increasing and sustained trend seen in the data (Fig. 5D), suggesting the new models are predictive and valid for further analysis.

Using the cell-type-specific models and simulation, we next interrogate the YAP-TFs switches in the new cell lines. Figure 5F, G display simulations of the YAP-SMAD and YAP-p73 complexes in response to increasing ITCH and RASSF1A. Interestingly, while RASSF1A/ITCH controls the inter-transition between the YAP complexes in a qualitatively similar manner across the cell types, simulation results indicate that the YAP switches display distinct quantitative behaviours regarding both the switch steepness and threshold. Specifically, the ITCH-mediated YAP switches are sigmoidal and most abrupt in MDA-MB-468, and the switching threshold is lowest in MDA-MB-231 (Fig. 5F). Moreover, the responses of the YAP complexes to RASSF1A are more linear and gradual in Hela and MDA-MB-231 cells compared to MDA-MB-468 (Fig. 5G).

Overall, these findings demonstrate the presence of the on/off-switches of YAP-TFs complexes coordinated by ITCH and RASSF1A in multiple cell types, but the specific pattern of the switches is dictated by the protein expression profile in the individual cell types. Importantly, our computational models provide an important tool that enables quantitative prediction of the switches’ context-specific behaviour.

### ITCH’s regulatory impact on YAP activities correlates with its essentiality for cell viability

Our analysis in the previous section demonstrates ITCH differentially regulates YAP transcriptional complexes depending on the cellular context. For example, ITCH potently controls YAP-SMAD/p73 formation in MDA-MB-468 and MCF-10A cells through steep molecular switches, while it does so in a more graded manner in MDA-MB-231 and Hela cells (Fig. 5D). Given that the formation of distinct YAP-TFs complexes induces differential transcriptional programs, we reasoned that the extent to which ITCH controls YAP complexes may be linked to its functional importance. To test this, we first assessed ITCH’s role in maintaining cell viability by obtaining its ‘essentiality score’ in 342 cancer cell lines from the DEPMAP data portal (29). By measuring the effect of systematic gene deletion on cell viability, DEPMAP stores essentiality scores (between -1 and 1) for thousands of genes in hundreds of cancer cell lines, where: -1 indicates strong essentiality (gene loss reduces cell viability), 0 indicates neutrality (gene loss has no change in cell viability), or 1 if the loss of gene increases cell viability instead.

To examine if the essentiality of ITCH is associated with its regulatory impact on YAP activities, we need to determine the latter in the corresponding cell lines. To this end, we obtained protein expression data of 375 human cancer cell lines from the Cancer Cell Line Encyclopedia (CCLE) consortium (47). Of these, 54 cell lines express detectable levels of the key model components (Fig. 6A) and thus were selected for further analysis (Fig. S6). Using the model customization as described in the previous section, we constructed 54 cell lines-specific models and used them to simulate the ITCH-mediated YAP switches in each cell line (Fig. 6B). Model simulations show that the YAP switch-like responses are a robust feature across many cell lines; however, the amplitude and threshold of the switches vary between the cells. To compute the impact of ITCH on YAP activities regulation, we defined and computed an *ITCH impact score* (given in Fig. 6A) for each cell line. The score quantifies the impact of variation of ITCH concentration on the changes in YAP-SMAD and YAP-p73 abundances. where a higher score indicates ITCH has stronger impact on YAP regulation.

**Figure 6.**
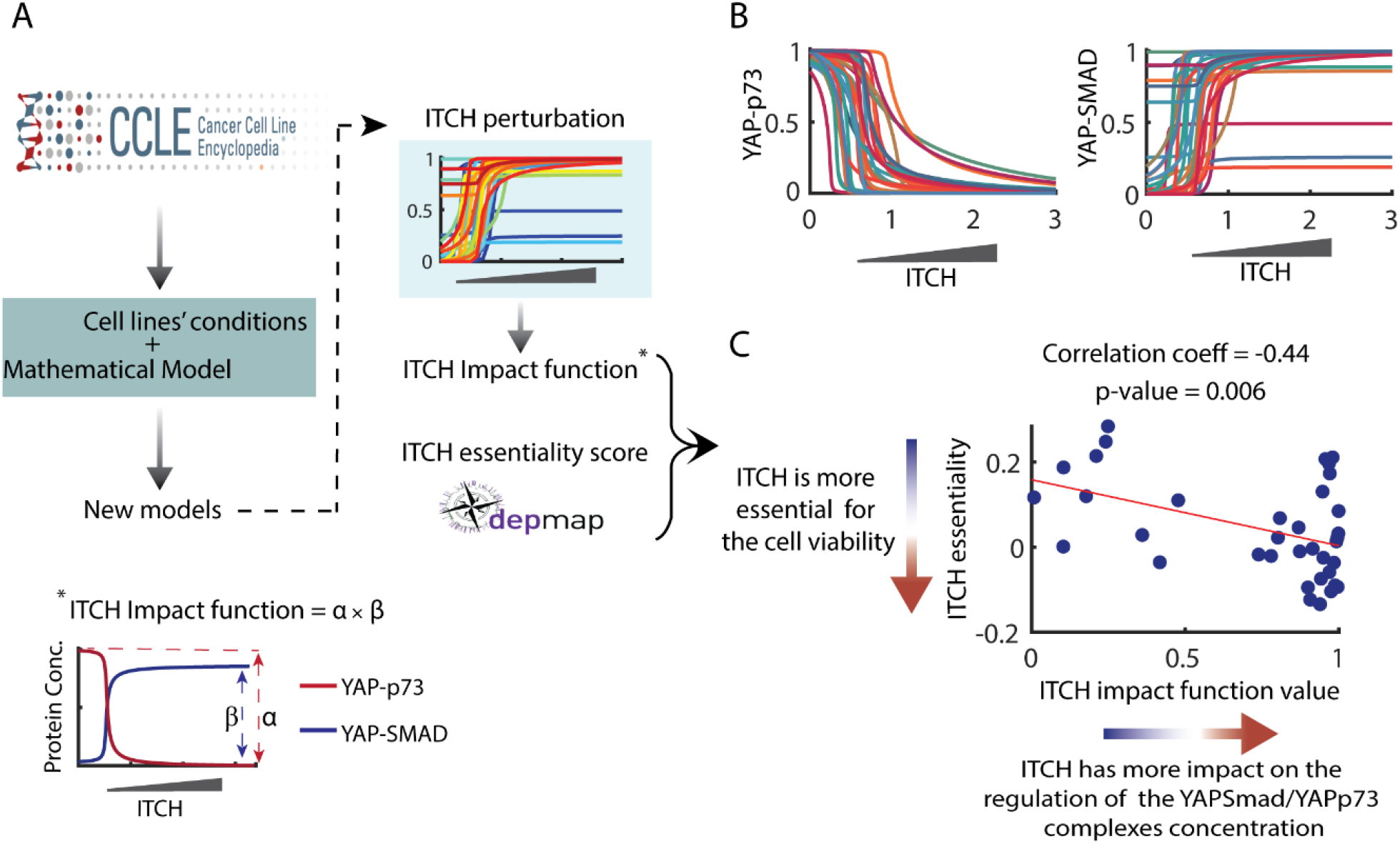
The impact of ITCH in the regulation of YAP activities is linked to its essentiality in cell viability. **(A)** A workflow used to simulate the ITCH-mediated YAP switches in multiple cancer cell lines using data from the CCLE database, and compute ITCH’s impact on YAP regulation. Briefly, CCLE database was used to find the initial condition of the components of model for 375 different cell lines. Among them 54 cell lines express detectable levels of the model components and thus were selected for further analysis. Using these initial conditions the existing model is adjusted to the 54 cell lines. *ITCH impact score* is defined and calculated for each cell line. ITCH essentiality score for the cell viability is also obtained for 342 cell lines using DEPMAP bio-portal. There are 36 cell lines that overlap between the DEPMAP and CCLE databases (i.e. having both essentiality and impact scores for ITCH). A more detailed description is given in Materials and Methods. **(B)** Simulation of the ITCH-dependent switches of YAP complexes in different cell lines using cell type specific models generated in (A). To perturb ITCH levels *in silico*, the total concentration of ITCH in each specific cell line model was changed from zero to three folds higher than the nominal values. Model simulations predict the presence of abrupt switch-like response of the YAP-TF complexes. **(C)** A significant and strong correlation is recorded between the ITCH impact function values and ITCH’s essentiality score across 36 different cell lines.

Since there are 36 cell lines that overlap between the DEPMAP and CCLE databases (i.e. having both essentiality and impact scores for ITCH), we performed correlation analysis between the two scores for these 36 cell lines. Strikingly, we observed a significant and strong negative correlation between the scores (correlation-coefficient= -0.44, p-value = 0.006), indicating that how essential YAP is in driving cell viability is positively associated with how potently it regulates the YAP transcriptional complexes. This finding provides a functional connection between the YAP-mediating role of ITCH and its biological importance.

## DISCUSSION

Mounting experimental evidence indicates that YAP can function both as a regulator of proliferative or apoptotic programs in cells. However, how YAP mediates its context-dependent function remains poorly understood. Upon activation, the TGF-β signalling pathway recruits YAP to induce oncogenic transcription through the formation of the YAP-SMAD complex. On the other hand, via RASSF1A the Hippo pathway triggers apoptotic transcription by promoting YAP-p73 binding. As these pathways display intimate crosstalk at multiple levels, it is likely that depending on the specific network state and condition YAP can favour assembly with a particular transcriptional factor to either accelerate or suppress tumour progression. Depsite this, how the opposing YAP transcriptional complexes are coordinated at the network-level has not been characterized. This is due largely to the paucity of integrative frameworks that can tease apart the network complexity. In this study, we developed new mathematical models describing the complex TGF-β/Hippo pathway crosstalk and undertook integrated modelling-experimental analyses to study the dynamic behaviour and regulation of the YAP-TFs complexes under various cellular contexts.

Several mathematical models were previously developed for the TGF-β or Hippo pathway separately, by us and others (1, 20, 32, 33). For example, Zi *et. al*. constructed an ODE model to investigate the effect of time- and dose-varying TGF-β stimulation on TGF-β signalling (33). Their analysis revealed a switch-like response of pSMAD2 to increasing TGF-β concentration at long-term stimulations. This observation is opposite to a linear short-term response of pSMAD2 to TGF-β and suggests the switch-like response may be critical for cell-fate determination. Nicklas *et. al*. developed a model considering phosphorylation of SMAD1/2, their binding to SMAD4 and shuttling of the complex to the nucleus (32). Using this model they investigated the underlying mechanisms of distinct dynamics of the TGF-β pathway under different network conditions. We have developed models of the Hippo pathway in crosstalk with other pathways including RAF-1/ERK and AKT, which have uncovered molecular switches that coordinate opposite cell fate decisions (1, 20). However, these models did not include YAP and only account for the signalling events upstream of YAP. A recent model considers crosstalk between TGF-β and YAP signalling (40), however, this model did not incorporate competing YAP complexes or the crosstalk mechanism mediated by RASSF1A/ITCH. Thus, the model in this study represents the first experimentally validated model describing in detail the mechanistic interplay between the TGF-β and Hippo-YAP pathways, which converge on the formation of transcriptionally opposing YAP-TF complexes.

Our integrative studies reveal for the first time the presence of molecular switches that regulate the competing formation of the YAP-SMAD and YAP-p73 complexes. As these complexes mediate opposing gene expression programs, the identified switches may provide a mechanism for cells to make a swift and robust decision toward distinct cellular fates (i.e. cell proliferation or death) that are induced by YAP. Indeed, molecular switches have been widely employed by living cells to control cell fate decision (48-50). These include bistable and sigmoidal switches that are brought about by positive feedback (51, 52) and non-feedback regulation, respectively (1, 53).

We found that RASSF1A and ITCH are critical coordinators of the YAP-TFs switches. A graded increase in RASSF1A expression can trigger an abrupt boost information of the apoptotic YAP-p73 complex, accompanied by an abrupt disassembly of the proliferative YAP-SMAD complex; while ITCH acts in the opposite manner. These findings are in line with the tumour-suppressive function of RASSF1A in various cancer types (54). Moreover, as YAP is competed for binding by SMAD and p73, the identified switches also demonstrate the threshold-controlled inter-transition of YAP between the complexes. For example, when ITCH expression exceeds a critical threshold, YAP flips from predominantly binding p73 to binding SMAD, thereby turning up proliferative gene expression while at the same time turning down apoptotic gene expression. Such coordinated switching between the functionally opposing complexes by YAP may further underpin the decisiveness of the cellular decision-making machinery.

Further, our model allowed us to perform model sensitivity analysis that systematically identifies the governing parameters that control the behaviour of the YAP switches. We found that the degradation rate of RASSF1A has the strongest influence on the switching threshold, while the dissociation rate of the YAP-p73 complex has the strongest impact on the steepness of the switches. These findings highlight the benefit of computational models as a valuable tool in teasing out the hidden factors regulating intricate dynamic behaviours within complex signalling networks, a task that would be more challenging using experimental approaches alone.

Signalling dynamics are strongly influenced by the cell-to-cell variability in protein expression (55). Although we have identified abrupt switches regulating the YAP complexes in MCF-10A cells, we wondered whether similar switches may also exist under other cellular contexts. To answer this, we characterized the protein expression of the key network nodes in additional breast cancer cell lines (MDA-MB-468, MDA-MB-231 and Hela) in comparison to MCF-10A cells, and utilised these data to generate new cell type-specific models based on the MCF-10A model. Simulation studies using these models demonstrate that the behaviour of the YAP switches is strongly determined by the specific expression profile of the network components in specific cell types. While the threshold-gated transition of YAP between the TF complexes is commonly observed, the degree of the switches (steepness) and the switching threshold vary depending on the cell types. For example, the switches appear most abrupt in MDA-MB-468 cells in response to increasing ITCH, while in Hela and MDA-MB-231 cells, ITCH perturbation results in relatively smoother changes in YAP-SMAD. The same pattern was also observed in response to RASSF1A perturbation. These results demonstrate the highly context-specific patterns of the switches regulating YAP transcriptional complexes and highlight the need to embrace network contexts when studying their behaviour.

Our analysis revealed the oncogenic role of ITCH by upregulating YAP-SMAD and suppressing YAP-p73 (Fig 3C, D). Considering the difference in amplitude of YAP response to ITCH perturbation in different cell lines (Fig. 5D) we wondered whether this amplitude is correlated with the overall impact of ITCH in cell viability. The importance of each gene in the cell viability is defined as genes essentiality score by DEPMAP Bio portal. In their study, each gene is knocked out in various cell lines and the essentiality score is calculated based on the effect of the gene knockdown on the changes in the cell viability. Therefore, an essentiality score was obtained for each gene in each cell line. Using this data we obtained the essentiality of ITCH in the viability of different cell lines. In the next step, we calculated the amplitude of YAP response to ITCH perturbation using the *ITCH impact function* (Fig. 6A). Interestingly, the results revealed a correlation between the significance of ITCH in the regulation of YAP activities and its essentiality score in cell lines (Fig. 6B). These results indicate the importance of the ITCH-mediated switch in YAP activities in the overall role of ITCH in cells.

Our studies implicate the E3 ligase ITCH as a novel regulator of the YAP-TFs complexes and YAP transcriptional activities. Specifically, ITCH stimulates proliferative target genes through enhancing YAP-SMAD while inhibiting YAP-p73 complex formation. This finding is consistent with published studies reporting oncogenic functions of ITCH in different malignancies (56-60). Overexpression of ITCH in breast cancer cells enhances epithelial to mesenchymal transition and its knockdown inhibits tumorigenicity and metastasis (31). Further, ITCH is significantly upregulated in invasive and metastatic breast cancer cases and its expression is associated with poorer survival (31). In addition, our analysis of pan-cancer patient data from The Cancer Genome Atlas (“https://www.cancer.gov/tcga.”) indicates ITCH is also upregulated in subsets of other cancer types including rectal adenocarcinoma, uterine carcinoma and colon adenocarcinoma (Fig. S6). Targeting ITCH may thus represent viable therapeutic strategies for the development of new treatment of specific cancers (30).

## MATERIAL AND METHODS

### Cell culture

Human mammary epithelial MCF-10A cells were cultured in basal Dulbecco’s Modified Eagle Medium (DMEM) (Gibco by Life Technologies) with 10% Fetal Bovine Serum (FBS) (Gibco by Life Technologies), and 1x GlutaMAX (Gibco by Life Technologies) at 37 °C and 5% CO2

MCF-10A cells obtained from the ATCC (USA) were cultured in high glucose Dulbecco’s Modified Eagle Medium (DMEM) (Gibco by Life Technologies) with 10% Fetal Bovine Serum (FBS) (Gibco by Life Technologies), and 1x GlutaMAX (Gibco by Life Technologies) at 37 °C and 10% CO2. Cells were used for experiments 3 days after initiation of differentiation. HeLa, MDA-MB-231 and MDA-MB-468 cell lines were obtained from the American Type Culture Collection (ATCC) and grown in the medium described above to culture MCF-10A cells. Cells were maintained in Minimum Essential Media (MEM) (Gibco by Life Technologies catalog #10370-021), with the addition of 10% (v/v) FBS, 1x GlutaMAX, and 1 mM sodium pyruvate (Gibco by Life Technologies) at 37 °C and 5% CO2.

### Plasmids, siRNA and transfection

MCF-10A cells were transfected with esiRNA at approximately 70–75% confluency using*Trans*IT-X2 (Mirus Bio) according to manufacturer’s instructions.

### Western Blotting

MCF-10A cells were serum-starved with DMEM containing 1× GlutaMAX and 0.2% BSA (w/v) for 24 h and exposed to TGFβ. Cells were lysed with RIPA lysis buffer supplemented with 10 μg/mL Aprotinin, 10 μg/mL Leupeptin, 1 mM PMSF and 1 mM sodium orthovanadate. Cells were then placed on ice, washed with cold PBS. Cells were scraped from the plate/well, and then cell suspension was collected and subjected to low-speed centrifugation (15,000 x g) for 15 minutes at 4oC. Pierce™ BCA Protein Assay Kit was used to quantify lysate’s protein concentration following the manufacturer’s protocol (61).

### Immunoblotting

Samples were boiled for 5 minutes at 96 °C and then resolved on SDS-PAGE gels (10% running and 4% stacking) at 80 V for 20 minutes, followed by 120 V for 1.5 hours in running buffer at 100 V for 1.5 hours or at 26 V overnight (for high molecular weight proteins). Transferred membranes were blocked with BSA blocking buffer on a rocking platform at room temperature for 1 hour. Membranes were then incubated in primary antibodies diluted in BSA blocking buffer overnight at 4°C. Membranes were washed three times, 5 minutes each, with 1X TBS-T while rocking. Secondary HRP-conjugated antibodies diluted in 5% (w/v) skim milk was added for 1 hour at room temperature then membranes were washed three times in 1X TBS-T. Membranes were immersed in Immobilon® Crescendo Western HRP Substrate (Merck) for 30 seconds, then ChemiDoc Imager (Bio-Rad Laboratories) was used to obtain immunoblot images following the manufacturer’s instruction (62). Densitometry analysis was performed on the detected bands using ImageLab, version 5.2.1 (Bio-Rad). β-actin or 14-3-3 were used as the loading controls. The intensity of each band was normalised against the intensity of their corresponding loading control. Bands correlating to phospho-proteins were further normalised to the bands of corresponding total protein levels.

The following antibodies were purchased from Cell Signaling Technology: pSMAD2 (S465) (3471), SMAD2 (86F7), YAP (14074), pYAP (Y357) (4911), pYAP (S127) (13008),, pSMAD3 (S423/S425). We purchased SMAD7 (ab216428) from abcam and RASSF1A(eB114-10H1) from e-bioscience.

### Mathematical modelling and simulations

In this paper, we constructed a model for the Hippo-TGF-β crosstalk. The biochemical reactions are described using ordinary differential equations (ODEs) known as chemical kinetic equations. The full reaction rates are provided in Supplementary Tables S1-2 and the best-fitted parameter sets used for simulations are given in Supplementary Information 1. The model construction and calibration processes were implemented in MATLAB (The MathWorks. Inc. 2020b), and the IQM toolbox.

### Model fitting and parameter estimation

To estimate the model parameters objective function *J* was used that quantifies the difference between the simulation results of the model and quantified experimental data:

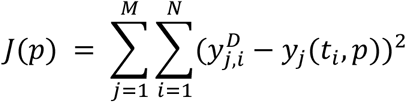

Here, *M* denotes the number of experimental data sets used for fitting and *N* is the number of time points in each set. *Y*_*J*_(*t*_*i*_, *p*) indicates simulation of component *j* at the time point *t*_*i*_ while parameter set *p* is used for the simulation. 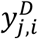 is the experimental data of component *j* at time point *t*_*i*_ and *w*_*j*_ is the weight of the component *j*.

The value of parameters was estimated to minimize the value of objective function *j*. Parameter estimation was undertaken using Global Optimization Toolbox and the *ga* function in MATLAB with the population size of 200 and generation number of 800.

Since it is likely to exist multiple optimal parameters sets that fit the experimental data (63), we repeated the GA procedure 500 independent times to recognise as many best-fitted parameter sets as possible for subsequent ensemble simulations. The selection criteria for choosing the best-fitted parameter sets was the objective function values for each of these sets must be under a cut-off threshold. As a result, we selected 11 best-fitted parameter sets for the model for further simulation and analysis, respectively. The code and parameter sets have been stored in Github(https://github.com/NguyenLab-IntegratedNetworkModeling/TGF_Hippo_crosstalk).

### Cell type specific modelling and simulation

The cells’ responses to external stimulations are heterogeneous. This diversity stem from variations in proteins concentration in different cells types. Therefore, to customize our calibrated model of MCF-10A cells to fit to Hela, MDA-MB-468 and MDA-MB-231 cancer cell lines, we used the relative expression data of proteins from western blot experiments (Fig 5A). The initial condition of the model was changed according to the relative proteins concentration to obtain three models specific to Hela, MDA-MB-468 and MDA-MB-231 cell lines. Therefore:

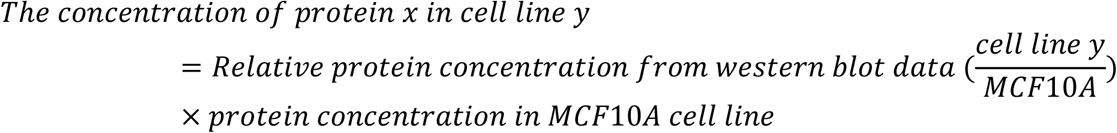

We also used relative protein expression of 375 human cancer cell lines from CCLE database to obtain customized models specific to these cell lines. Among the 375 cell lines only 54 of them expressed a detectable amount of key component proteins and therefore considered for model customization. As described above, this data was used to modify the initial condition of proteins in the calibrated model to fit the other cell lines condition. MDA-MB-468 is one of the 375 cell lines in CCLE database and therefore we used MDA-MB-468 customized model obtained from the previous section as a reference model to change its initial condition to generate models for the other 53 cell lines.

## SUPPLEMENTARY FIGURES

**Figure S1.**
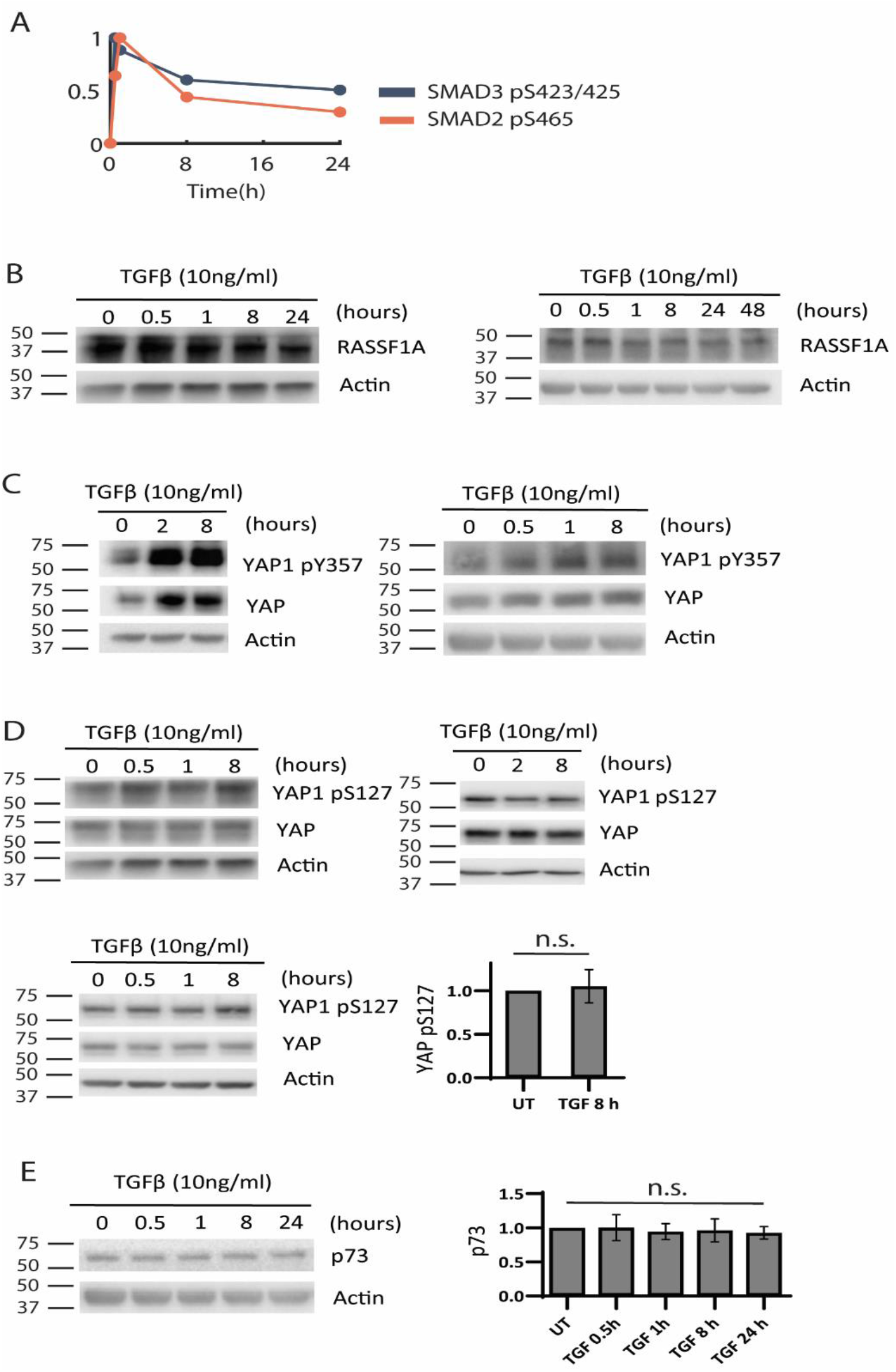
Characterization of the dynamic TGF-β-Hippo pathway signalling crosstalk. **(A)** Quantification of the western blot data in Figure 1A, using ImageJ. **(B-E)** Time-course western blot analysis and associated quantification of various signalling proteins in response to TGF-β stimulation (10ng/ml) treatment in MCF-10A cells. Cells lysates were immunoblotted using the antibodies as specified with actin as a loading control. For (B-D), different biological replicates of the time-course profiling experiments are displayed.

**Figure S2.**
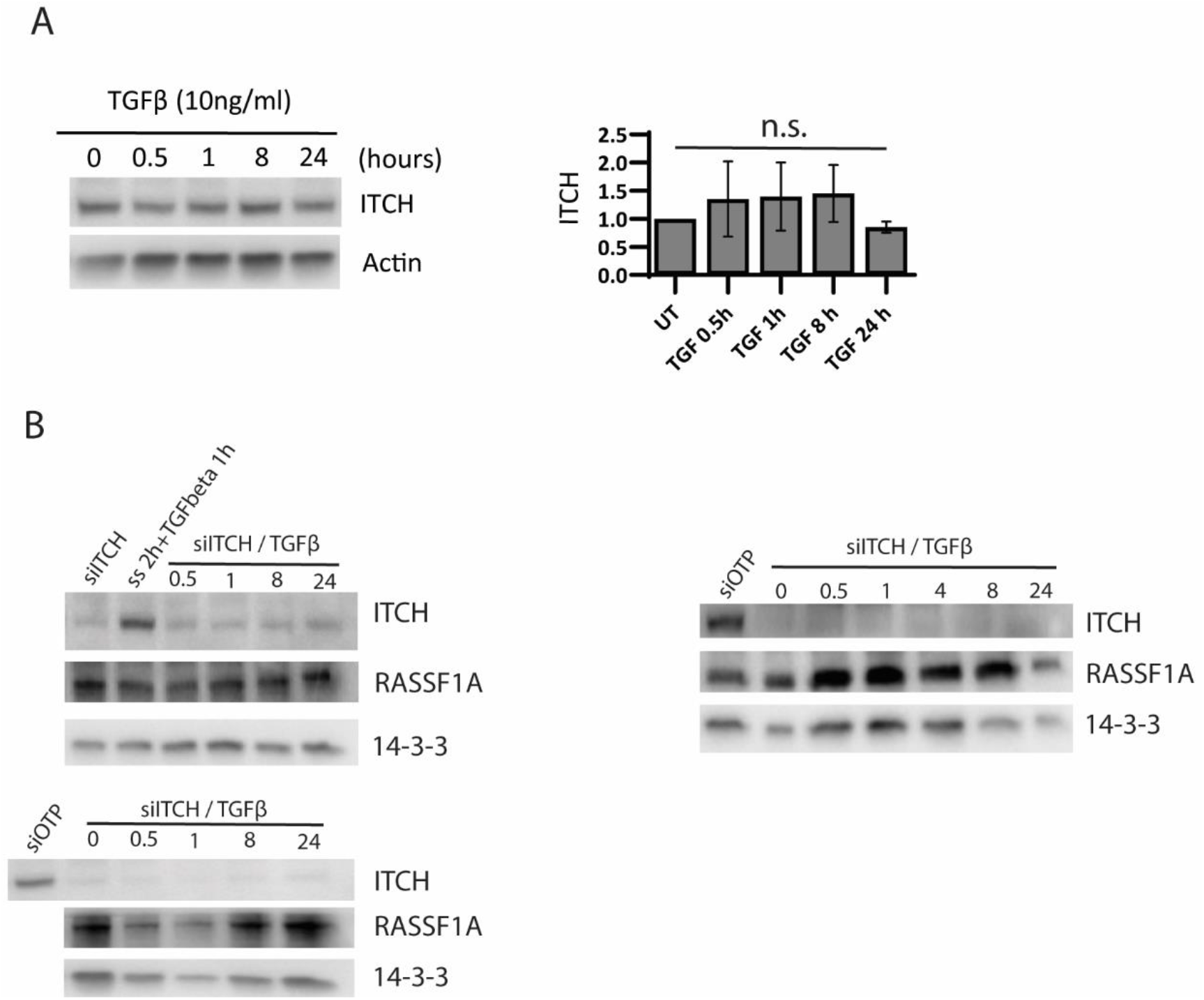
Impact of TGF-β stimulation on the components concentration. **(A)** TGF-β stimulation has no significant effect on ITCH concentration over time. **(B)** The effect of TGF-β stimulation on RASSF1A concentration in ITCH knocked down MCF-10A cells (Figure 2D demonestrates the quantified results). RASSF1A level remained unchanged in response to TGF-β stimulation in ITCH knocked down MCF-10A cells.

**Figure S3.**
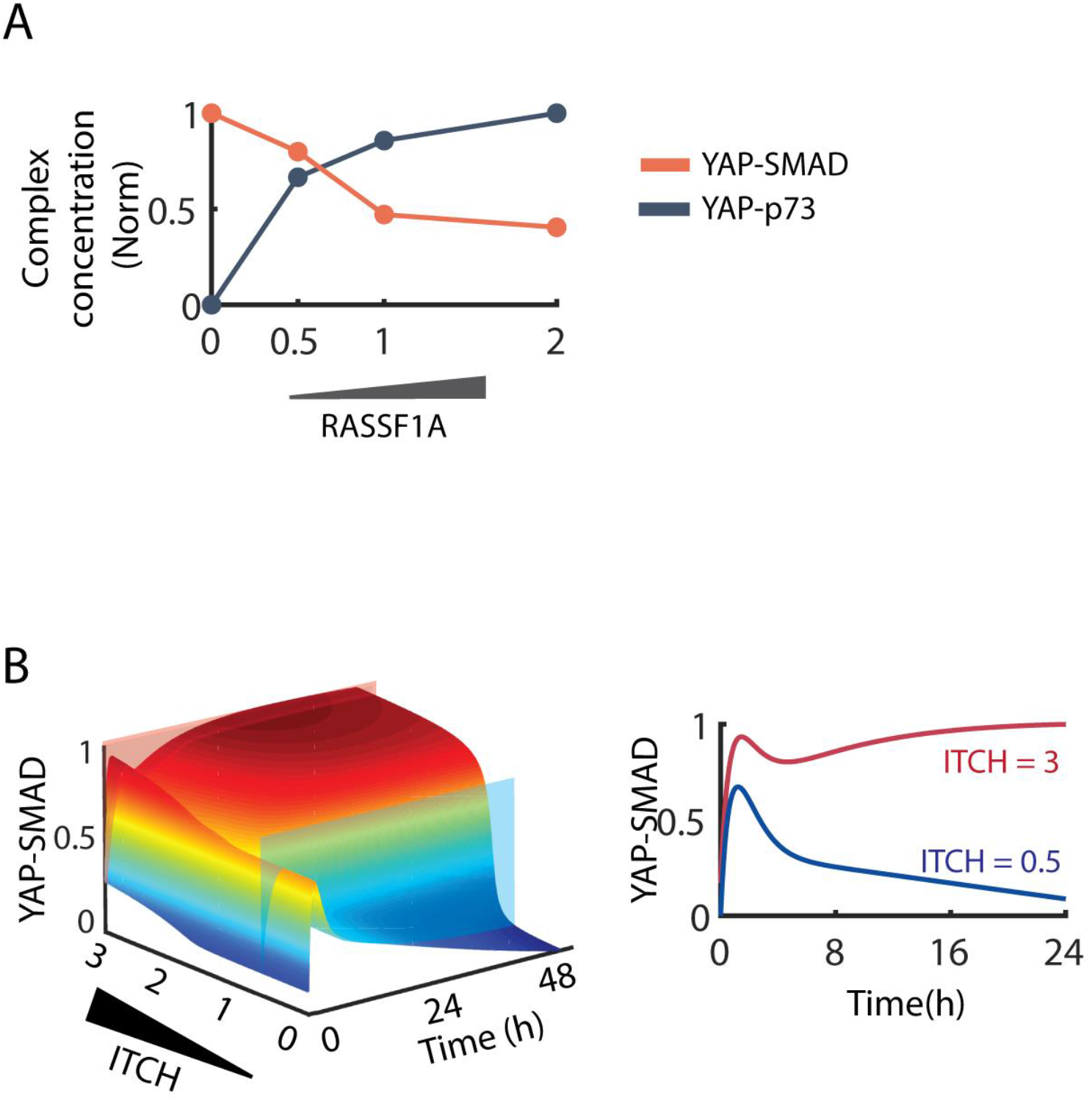
**(A)** Quantitative data of YAP-SMAD and YAP-p73 concentration at different RASSF1A expression levels in U2OS inducible cell lines, from Pefani *et al* (3) **(B)** 3-D simulations of the YAP-SMAD and YAP-p73 complexes in response to increasing ITCH concentration. At the low level of ITCH YAP-SMAD displays an overshoot dynamics after TGFβ stimulation. High concentration of ITCH leads to a sustained activation of YAP-SMAD after TGFβ treatment.

**Figure S4.**
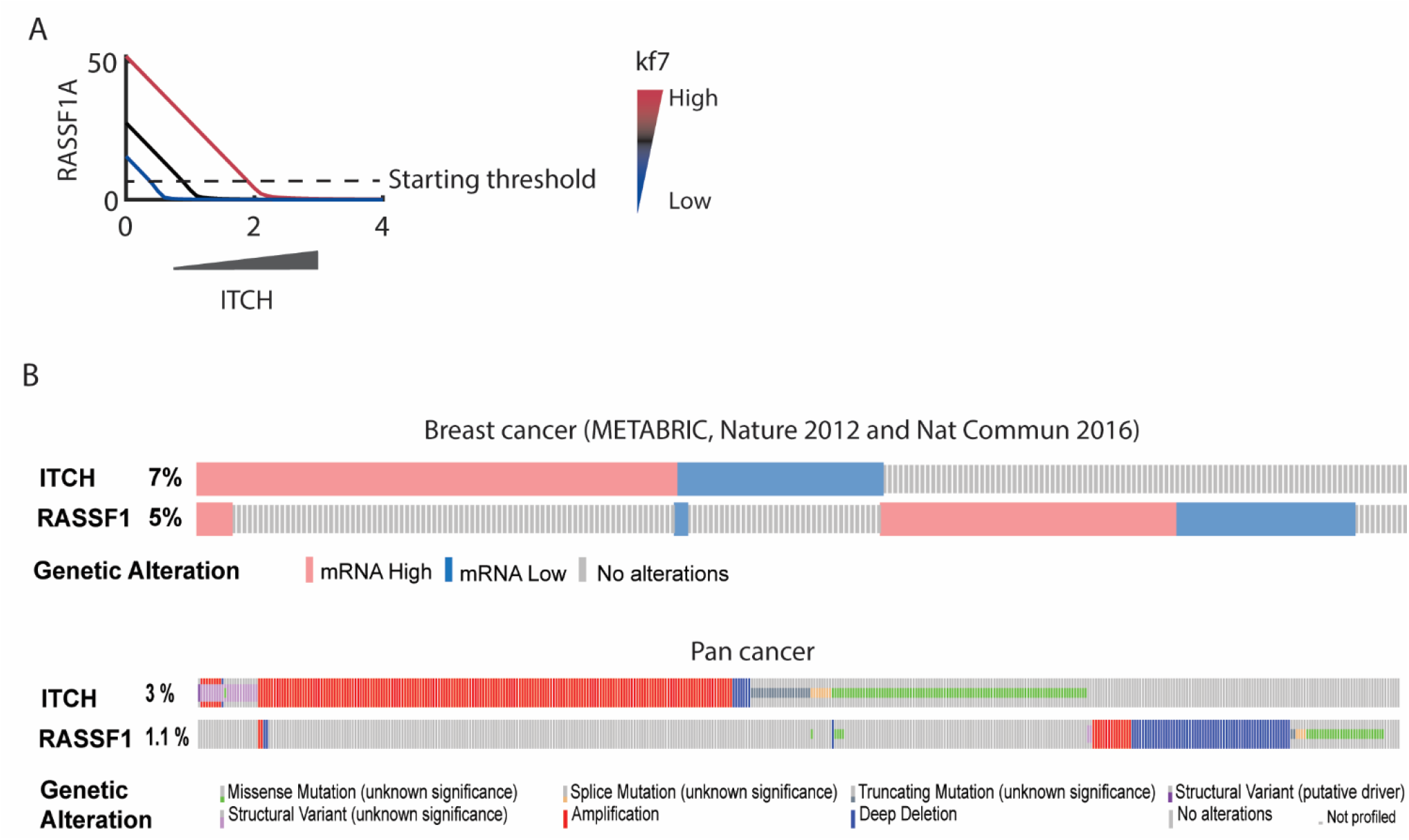
(A) Increasing RASSF1A’s synthesis rate needs higher ITCH concentrations to degrade RASSF1A and trigger the switching between YAP-SMAD and YAP-p73. **(B)** Analysing the co-expression profiles of RASSF1 and ITCH across large cohorts of breast and pan-cancer patients (64) shows that patients with down-regulated RASSF1 rarely exhibit overexpression of ITCH compared to those with normal or upregulated RASSF1.

**Figure S5.**
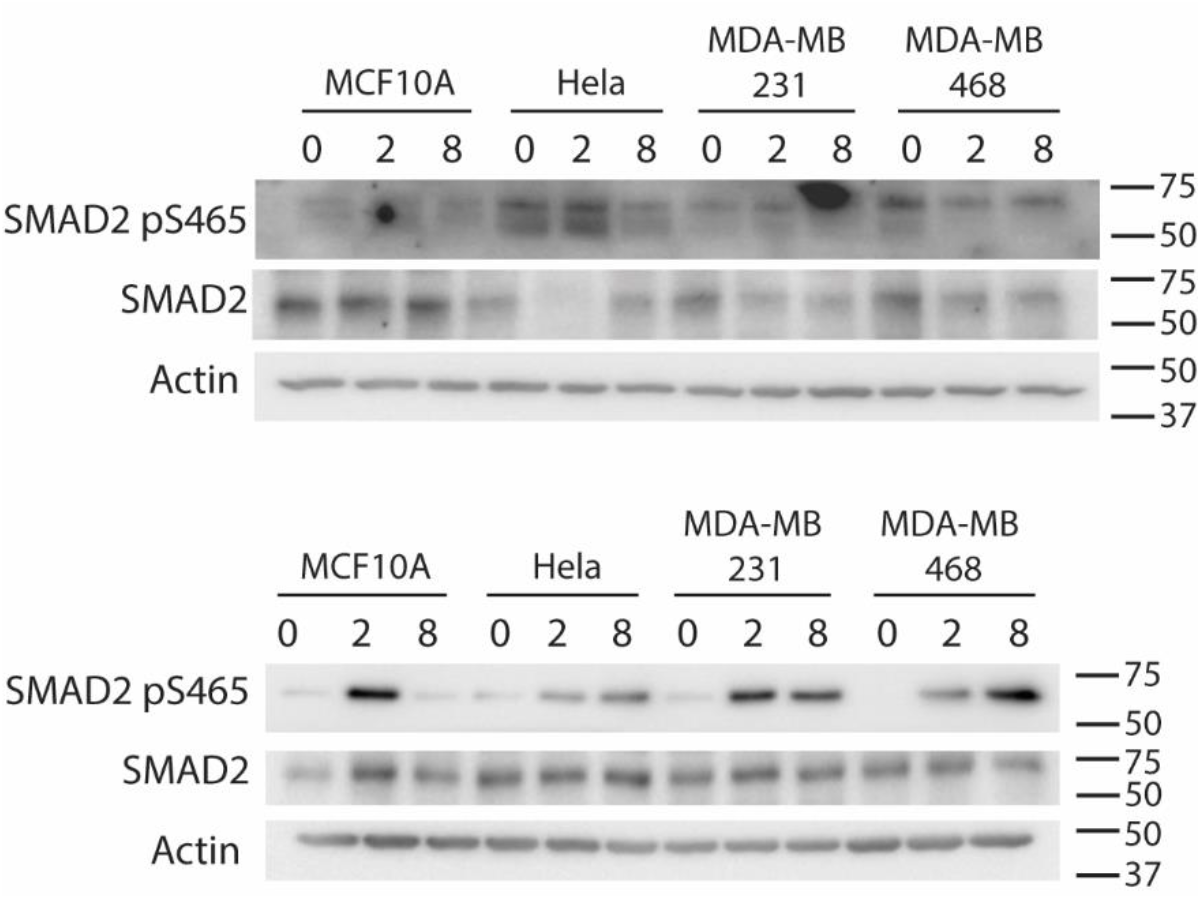
Analysis of pSMAD2 dynamics in response to TGF-β stimulation in four different cell lines. Investigating pSMAD2 dynamics in response to 8 hours of TGF-β (10ng/ml) across MCF-10A and Hela, MDA-MB-468 and MDA-MB-231 using western blot (second and third replicates). Cells lysates were immunoblotted using the antibodies as specified with actin as a loading control.

**Figure S6.**
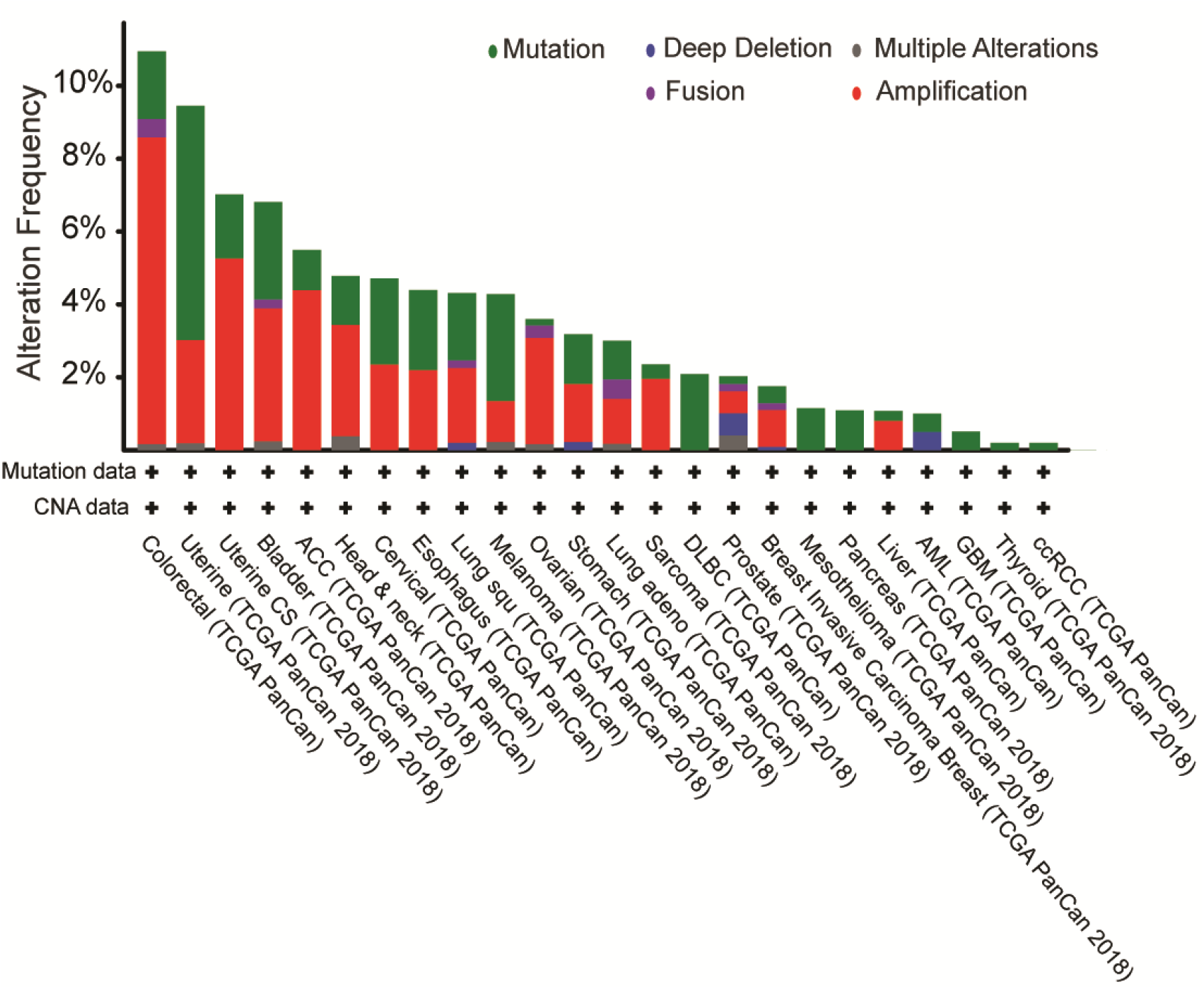
Alteration frequency of the ITCH gene in cancer patients analyzed from TCGA database using cBioPortal (www.cbioportal.org)(64).

## SUPPLEMENTARY TABLES

**Table S1.** Reactions and rate equations of the TGFβ-Hippo crosstalk model.

|  | Reactions | Reaction rates |
| --- | --- | --- |
| R1 | ↔ Receptor | $kf1 - (kf1/13.9) * \text{Receptor}$ |
| R2 | $\text{Receptor} \leftrightarrow \text{TGFb\_Rec}$ | $kf2 * \text{Receptor} * (\text{TGF} + \text{TGF0}) - kr2 * \text{TGFb\_Rec}$ |
| R4 | $\text{SMAD2} \leftrightarrow \text{pSMAD2}$ | $kf3 * \text{TGFb\_Rec} * \text{SMAD2} / (\text{km3} + \text{SMAD2}) - kr3 * \text{pSMAD2}$ |
| R6 | $\text{Receptor} \leftrightarrow$ | $kf4 * \text{Smad7} * \text{TGFb\_Rec} / (\text{km4} + \text{TGFb\_Rec})$ |
| R7 | $\text{c-Abl} \leftrightarrow \text{pc-Abl}$ | $kf5 * \text{TGFb\_Rec} * \text{c-Abl} / (\text{km5} + \text{c-Abl}) - kr5 * \text{pc-Abl}$ |
| R8 | $\text{Cyto\_ITCH} \leftrightarrow \text{pm\_ITCH}$ | $kf6 * \text{TGFb\_Rec} * \text{Cyto\_ITCH} / (\text{km6} + \text{Cyto\_ITCH}) - kr6 * \text{pm\_ITCH}$ |
| R9 | $\leftrightarrow \text{Cyto\_RASSF1A}$ | $kf7 - kr7 * \text{Cyto\_RASSF1A}$ |
| R10 | $\text{Cyto\_RASSF1A} \leftrightarrow \text{pm\_RASSF1A}$ | $kf8 * \text{TGFb\_Rec} * \text{Cyto\_RASSF1A} / (\text{km8} + \text{Cyto\_RASSF1A}) - kr8 * \text{pm\_RASSF1A}$ |
| R11 | $\text{pm\_RASSF1A} \leftrightarrow$ | $kf9 * (\text{pmITCH}) * \text{pm\_RASSF1A} / (\text{km9} + \text{pm\_RASSF1A})$ |
| R12 | $\text{YAP} + \text{pSMAD2} \leftrightarrow \text{YAP-SMAD}$ | $kf11 * \text{YAP} * \text{pSMAD2} - kr11 * \text{YAP-SMAD}$ |
| R13 | $\text{YAP} \leftrightarrow \text{pYAP\_y357}$ | $kf12 * \text{pc-Abl} * \text{YAP} / (\text{km12} + \text{YAP}) - kr12 * \text{pYAP\_y357}$ |
| R14 | $\text{P73} + \text{pYAP\_y357} \leftrightarrow \text{pYAP357-p73}$ | $kf13a * p73 * \text{pYAP\_y357} * \text{Cyto\_RASSF1A} / (\text{km131} + \text{Cyto\_RASSF1A}) + kf13b * p73 * \text{pYAP\_y357} * \text{pm\_RASSF1A} / (\text{km132} + \text{pm\_RASSF1A}) - kr13 * \text{pYAP357-p73}$ |
| R15 | $\text{P73} + \text{pYAP} \leftrightarrow \text{YAP-p73}$ | $kf14a * p73 * \text{pYAP\_y357} * \text{Cyto\_RASSF1A} / (\text{km141} + \text{Cyto\_RASSF1A}) + kf14b * p73 * \text{pYAP\_y357} * \text{pm\_RASSF1A} / (\text{km142} + \text{pm\_RASSF1A}) - kr14 * \text{YAP-p73}$ |
| R16 | $\leftrightarrow \text{SMAD7}$ | $kc151 * \text{YAP-SMAD} / (\text{km151} + \text{YAP-SMAD}) + kc152 * \text{pSMAD2} / (\text{km152} + \text{pSMAD2}) - (kr15) * \text{SMAD7}$ |

**Table S2.** Ordinary differential equations of the TGFβ-Hippo crosstalk model. The reaction rates are given in Table S1. The initial conditions are representative values.

| Left hand Sides | Right hand Sides | Initial Conditions (nM) |
| --- | --- | --- |
| d[Receptor]/dt | +R1-R2 | 13.9 |
| d[TGFb_Rec]/dt | +R2-R4 | 0 |
| d[SMAD2]/dt | -R3 | 107 |
| d[pSMAD2]/dt | +R3-R11 | 0 |
| d[c-Abl]/dt | -R5 | 1 |
| d[pc-Abl]/dt | +R5 | 0 |
| d[Cyto_ITCH]/dt | -R6 | 290 |
| d[pm_ITCH]/dt | +R6 | 0 |
| d[Cyto_RASSF1A]/dt | +R7-R8 | 10 |
| d[pm_RASSF1A]/dt | +R8-R9 | 0 |
| d[YAP]/dt | -R11-R12-R14 | 178 |
| d[YAP-SMAD]/dt | +R11 | 0 |
| d[p73]/dt | -R13-R14 | 20 |
| d[pYAP_y357]/dt | +R12-R13 | 0 |
| d[pYAP357-p73]/dt | +R13 | 0 |
| d[YAP-p73]/dt | +R14 | 0 |
| d[SMAD7]/dt | +R15 | 0 |

**Table S3.** Representative best-fitted parameter set. All the best-fitted parameters are provided in a separated Supplementary Table.

| **Parameter** | **Value** | **Unit** | **Parameter** | **Value** | **Unit** |
| --- | --- | --- | --- | --- | --- |
| 'kf1' | 0.613762 | nM sec<sup>-1</sup> | 'kr11' | 0.234963 | sec<sup>-1</sup> |
| 'kf2' | 141.5794 | nM<sup>-1</sup> sec<sup>-1</sup> | 'kf12' | 141.9058 | sec<sup>-1</sup> |
| 'kr2' | 279.8981 | sec<sup>-1</sup> | 'kr12' | 1.114295 | sec<sup>-1</sup> |
| 'kf3' | 35399.73 | sec<sup>-1</sup> | 'km12' | 0.005715 | nM |
| 'km3' | 99311.61 | nM | 'kf13a' | 73960.53 | nM<sup>-1</sup> sec<sup>-1</sup> |
| 'kr3' | 77446.18 | nM<sup>-1</sup> | 'kf13b' | 0.43451 | nM<sup>-1</sup> sec<sup>-1</sup> |
| 'kf4' | 95279.62 | sec<sup>-1</sup> | 'km131' | 16143.59 | nM |
| 'km4' | 77268.06 | nM | 'km132' | 95060.48 | nM |
| 'kf5' | 17782.79 | sec<sup>-1</sup> | 'kr13' | 3097.419 | sec<sup>-1</sup> |
| 'km5' | 5046.613 | nM | 'kf14a' | 165.1962 | nM<sup>-1</sup> sec<sup>-1</sup> |
| 'kr5' | 38.54784 | sec<sup>-1</sup> | 'kf14b' | 0.017989 | nM<sup>-1</sup> sec<sup>-1</sup> |
| 'kf6' | 81470.43 | sec<sup>-1</sup> | 'kr14' | 0.12218 | nM |
| 'km6' | 6870.684 | nM | 'km141' | 3639.15 | nM |
| 'kr6' | 28.84032 | sec<sup>-1</sup> | 'km142' | 55207.74 | sec<sup>-1</sup> |
| 'kf7' | 25.64484 | nM sec<sup>-1</sup> | 'kc151' | 1.69E-05 | nM sec<sup>-1</sup> |
| 'kr7' | 1.064143 | sec<sup>-1</sup> | 'kc152' | 1.69E-07 | nM sec<sup>-1</sup> |
| 'kf8' | 33.03695 | sec<sup>-1</sup> | 'km151' | 97949 | nM |
| 'km8' | 0.145546 | nM | 'km152' | 0.059979 | nM |
| 'kr8' | 427.5629 | sec<sup>-1</sup> | 'kr15' | 0.001052 | sec<sup>-1</sup> |
| 'kf9' | 0.096383 | sec<sup>-1</sup> | **TGFB0** | 1 | nM |
| 'km9' | 0.184077 | nM | **TGFB** | 100 | nM |
| 'kf11' | 100.6932 | nM<sup>-1</sup> sec<sup>-1</sup> |  |  |  |

